# Development of a joint evolutionary model for the genome and the epigenome

**DOI:** 10.1101/293076

**Authors:** Jia Lu, Xiaoyi Cao, Sheng Zhong

## Abstract

**Background:** Interspecies epigenome comparisons yielded functional information that cannot be revealed by genome comparison alone, begging for theoretical advances that enable principled analysis approaches. Whereas probabilistic genome evolution models provided theoretical foundation to comparative genomics studies, it remains challenging to extend DNA evolution models to epigenomes.

**Results:** We present an effort to develop *ab initio* evolution models for epigenomes, by explicitly expressing the joint probability of multispecies DNA sequences and histone modifications on homologous genomic regions. This joint probability is modeled as a mixture of four components representing four evolutionary hypotheses, namely dependence and independence of interspecies epigenomic variations to sequence mutations and to sequence insertions and deletions (indels). For model fitting, we implemented a maximum likelihood method by coupling downhill simplex algorithm with dynamic programming. Based on likelihood comparisons, the model can be used to infer whether interspecies epigenomic variations depend on mutation or indels in local genomic sequences. We applied this model to analyze DNase hypersensitive regions and spermatid H3K4me3 ChIP-seq data from human and rhesus macaque. Approximately 5.5% of homologous regions in the genomes exhibited H3K4me3 modification in either species, among which approximately 67% homologous regions exhibited sequence-dependent interspecies H3K4me3 variations. Mutations accounted for less sequence-dependent H3K4me3 variations than indels. Among transposon-mediated indels, ERV1 insertions and L1 insertions were most strongly associated with H3K4me3 gains and losses, respectively.

**Conclusion:** This work initiates a class of probabilistic evolution models that jointly model the genomes and the epigenomes, thus helps to bring evolutionary principles to comparative epigenomic studies.

## Background

Milestones of mathematical modeling of DNA evolution were marked by base substitution models in early 1980s [1–3], extension to incorporation of sequence insertions and deletions (indels) in early 1990s [4], and differential treatments of cis-regulatory sequences in the 2000-2010s [5–12]. The rise of interspecies transcriptome comparisons in 2000s [13–16] inspired a series of transcriptome comparison models and evolution models [17–19]. Benefits of joint analysis of interspecies variations of genomes and transcriptomes [20] demanded and eventually led to development of a joint probabilistic evolution model of the genome and the transcriptome [21].

Interspecies epigenome comparisons facilitated discoveries of functions of genomic sequences [22–26]. However, analyses of epigenome evolution remain observational, leading to divergent opinions on the dependence of epigenome conservation on sequence conservation. Some studies reported correlations between genomic and epigenomic changes [27, 28], whereas other studies revealed poor sequence conservation in homologous regions demarcated with the same histone modifications [29–31]. In much shorter timescale, sequence independent passage of histone modifications was observed in multiple generations [32, 33]. The development of evolutionary models for epigenomes would bring mathematical rigor to comparative epigenomics, and provide a model competition framework for evaluation of different hypotheses.

In this manuscript, we describe an effort on derivation of the joint probability of a pair of homologous genomic sequences and histone modifications on these sequences. We started with considering four hypotheses, where interspecies epigenomic variations (1) depend only on sequence mutations, or (2) depend only on sequence indels, or (3) depend on both mutations and indels, or (4) are independent of sequence mutations and indels. We formulated each hypothesis into a probabilistic evolution model, and developed a likelihood competition approach for model selection. This model competition approach enabled systematic evaluation of the four evolutionary hypotheses on any homologous sequences.

## Results

Our goal is to develop a probabilistic evolutionary model for a pair of homologous genomic regions that include the genomic sequences and histone modifications. If we denote the pair of homologous genomic regions as *A* and *B*, our goal is to derive the joint probability *P*(*A*, *B*). For this purpose, we introduce the following notations, model assumptions, and alternative hypotheses on evolution of genome and histone modifications.

### Notations

We introduce three sets of notations, including indices, observed data, and model parameters. The indices are *h* for indexing histone modifications (*h* = {1, 2, …, *H*}), *m* and *n* for indexing nucleotide positions in two DNA sequences, respectively, and *k* for indexing nucleotide positions in a pair of aligned sequences.

The observed data are denoted as follows. *A*^0^, *B*^0^ denote a pair of homologous genomic sequences. *A*^*h*^, *B*^*h*^ denote the states of the *h*^th^ histone modification on *A*^0^, *B*^0^. *A*, *B* denote a pair of homologous regions, including the homologous genomic sequences and the states of each histone modification on these sequences, where *A* = {*A*^0^, *A*^1^, …, *A*^*H*^}, and *B* = {*B*^0^, *B*^1^, …, *B*^*H*^}. Let *s*_*A*_ and *s*_*B*_ denote the lengths of *A*^0^ and *B*^0^. Let *a*_0,*m*_ and *b*_0,*n*_ denote the *m*^th^ and the *n*^th^ bases of sequences *A*^0^ and *B*^0^, where *a*_0,*m*_, *b*_0,*n*_ = {*A*, *C*, *G*, *T*}. Let *a*_*h*,*m*_ and *b*_*h*,*n*_ denote the states of the *h*^th^ histone modification at positions *m* and *n* in *A*^*h*^, *B*^*h*^, where *a*_*h*,*m*_ = {0,1} and *b*_*h*,*n*_ = {0,1}. Let 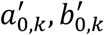 denote the nucleotides or indels on the *k*^th^ position of an aligned pair of sequences, where 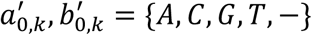. Let 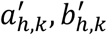 denote the states of the *h*^th^ histone modification on the *k*^th^ position in a pair of aligned sequences, where 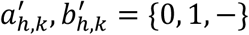. Finally, we denote an alignment of two sequences as a *path*, that is 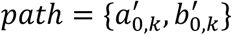.

The model parameters include *π*_Α_, *π*_C_, *π*_T_, *π*_G_, denoting the equilibrium probabilities of the four nucleotide bases. Let 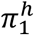 denote the global equilibrium probability, that is the equilibrium probability of having the *h*^th^ histone modification on any genomic location, and 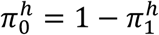. Let 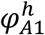 denote the local equilibrium probability, that is the probability of having the *h*^th^ histone modification on genomic region *A*, and 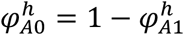. Denote sequence deletion rate as *μ*, insertion rate as *λ*, and substitution rate as *s*. Let *κ*^*h*^ be the rate of switch between 0 and 1, that is installing (0 to 1) or removing (1 to 0) for the *h*^th^ histone modification. Let *t* denote evolutionary time.

### Model assumptions

We assume that the state for each histone modification on each genomic location is binary, that is *A*^h^ and *B*^h^ are sequences of 0’s and 1’s with the same lengths as *A*^0^ and *B*^0^ (Figure 1). For example, a 5nt sequence of ACGTA (*A*^0^ = ACGTA) that is within an H3K9me3 peak (denote *A*^*h*=*H*3*K*9*me*3^ as *A*^1^) can be written as 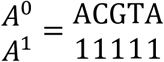. For another example, a 10nt sequence ACGTAGGGGG (*B*^0^ = ACGTAGGGGG) with the first 5 bases covered by an H3K9me3 peak and the second 5 bases not covered by any H3K9me3 peak can be written as 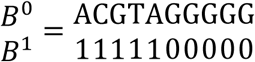, where *B*^1^ denotes the states of H3K9me3. Our second assumption is the widely adopted Pulley principle, namely that genomic evolutionary processes are reversible [3]. Our third assumption is that evolutionary changes of the DNA do not depend on evolutionary changes of epigenome.

**Figure 1.**
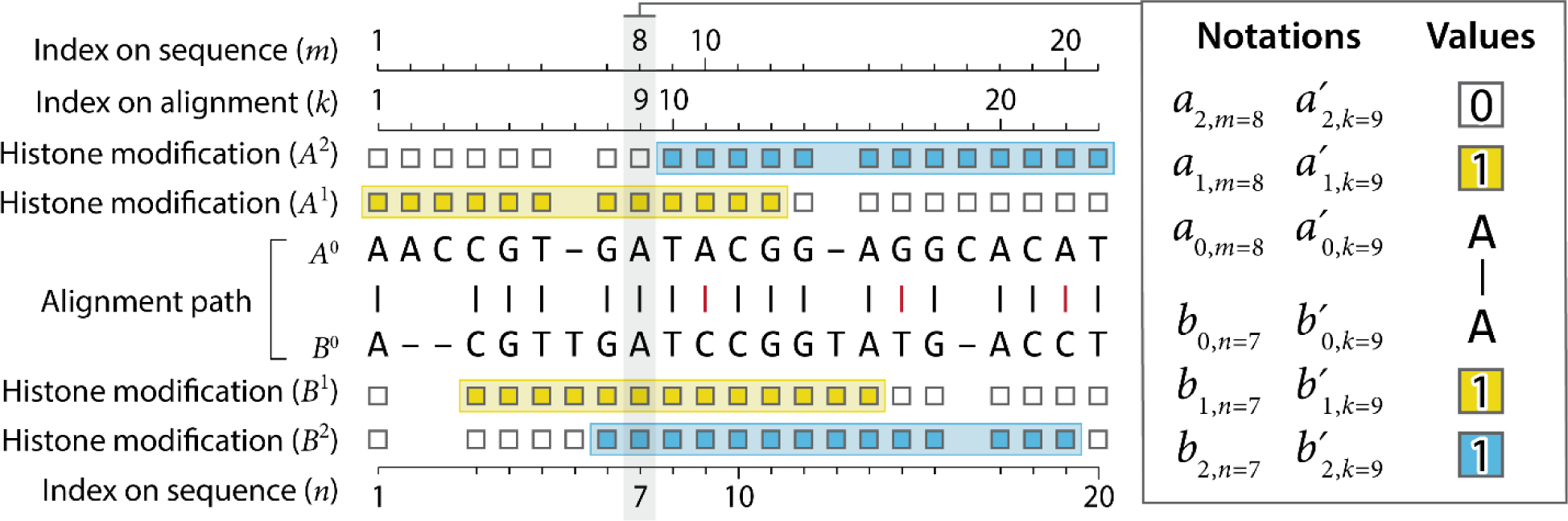
Data types and annotations. A pair of homologous sequences (*A*^0^, *B*^0^) are aligned, where −, | (black) and | (red) are indels, matches and mismatches, respectively. Base locations on each original sequence are indexed by *m* and *n* (indices on sequence). Base locations after sequence alignment are indexed by *k* (indices on path). Peak regions of two histone modifications *A*^1^, *B*^1^, and *A*^2^, *B*^2^ are shown as yellow and blue bands, respectively. A given histone modification on a given sequence, for example *A*^1^, is recorded by binary values on each base, with 1 being inside a peak and 0 being outside the peaks. Insert: notations and values of a specific position. On the 8^th^ position of sequence *A*^0^, the base is A (*a*_0,*m*=8_ = A). This base becomes the 9^th^ base after alignment 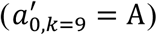. This base is inside a peak of the first (yellow) histone modification (*a*_*h*=1,*m*=8_ = 1) but outside any blue peaks (*a*_*h*=2,*m*=8_ = 0). If we use the base index after sequence alignment (*k*), that include indels, the above notations and values become 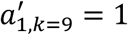 and 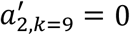.

### Development of a probabilistic framework for epigenome evolution

With the above introduced notations, our goal is to derive *P*(*A*, *B*) = *P*(*A*^0^, *A*^1^, …, *A*^*H*^, *B*^0^, *B*^1^, …, *B*^*H*^), where *A*^0^, *B*^0^ are homologous genomic sequences and *A*^*h*^, *B*^*h*^ (*h* = {1, …, *H*}) are histone modifications on *A*^0^, *B*^0^. To specify such a joint probability, we considered two types of dependency structures. First, descendent genomic sequence depends on ancestral sequence, and histone modifications depend on their underlying genomic sequence. The challenge of using such a dependency structure lies in the lack of complete knowledge of how genomic sequence determines the histone modifications, and therefore generally speaking *P*(*A*^*h*^|*A*^0^) cannot be specified. In the second type of dependency structure, descendent genomic sequence depends on the ancestral sequence, and histone modifications on the descent sequence depend on the histone modifications on the ancestral sequence. Furthermore, the evolutionary changes of each type of histone modification may depend on the genomic sequence changes (Figure 2A) or not (Figure 2B), and conditional on sequence changes the evolutionary changes of different histone modifications are independent of each other (conditional independence) (see Discussion). We elected to specify the joint probabilities with the second type of dependency structure.

**Figure 2.**
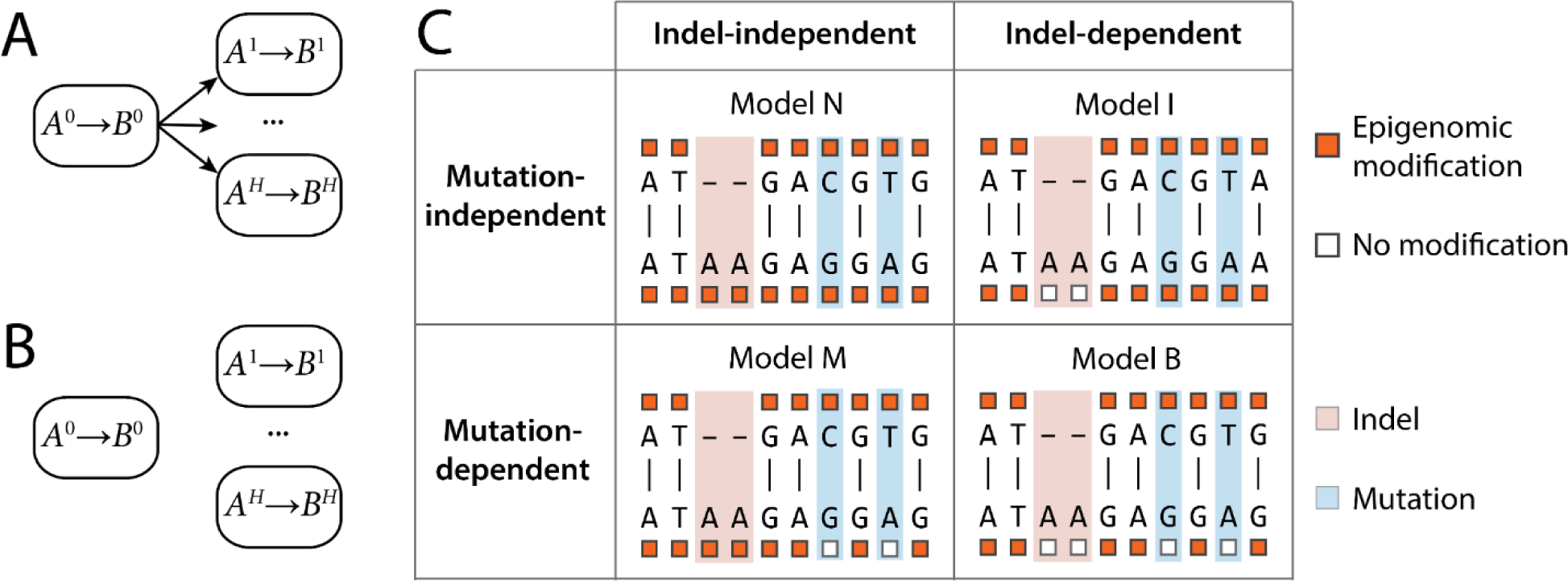
Dependency structures reflecting different evolutionary hypotheses. (A) Interspecies changes of the *h*^th^ histone modification (*A*^*h*^ → *B*^*h*^) depend on genomic sequence changes (*A*^0^ → *B*^0^). (B) Interspecies epigenomic changes do not depend on sequence changes. (C) A 2×2 table summarizing assumed dependencies to specific types of sequence changes in each model. Upper (bottom) row: models assuming independence (dependence) of sequence mutations. Left (right) column: models assuming independence (dependence) of sequence indels.

Based on the second type of dependency structure, we have

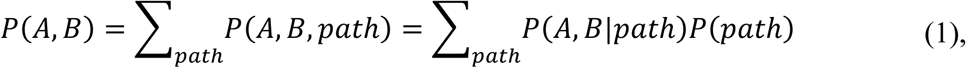

where *path* is an evolutionary path of homologous sequences, corresponding to an alignment of *A*^0^ and *B*^0^ (Figure 1). Any probabilistic expression of sequence alignment can be used for *P*(*path*), and in the work we employ the widely adopted TKF model as the analytical form of *P*(*path*) [4]. *P*(*A*, *B*|*path*) is the probability of observing a pair of homologous sequences and their epigenomes conditional on the sequence alignment. Because all sequence information is contained in *path*, due to conditional independence, we have:

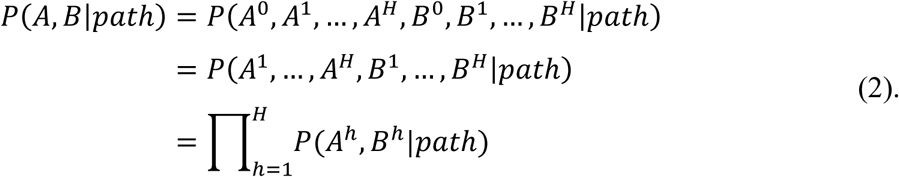

Applying previously introduced notations, we have:

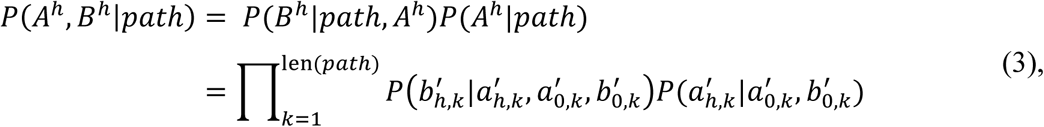

where len(*path*) is the length of the aligned sequence pair *A*^0^, *B*^0^ (first lane, Figure 1). Taking Equations (1) – (3) together, we have obtained a probabilistic statement of the observing a pair of homologous sequences and their respective histone modifications. Hereafter, we call Equations (1) – (3) the LCZ model. The LCZ model is fully specified when 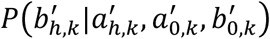 and 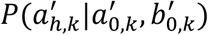 are specified.

### Translation of alternative evolutionary hypotheses into probabilistic models

We restricted this work to considerations of two types of sequence changes, namely mutations and indels. A total of four possible evolutionary hypotheses can be posed, that are (1) epigenome changes are independent of sequence changes (Model N), (2) epigenome changes depend on sequence mutations but are independent of sequence indels (Model M), (3) epigenome changes depend on sequence indels but not sequence mutations (Model I), and (4) epigenome depend on both mutations and indels (Model B, Figure 2C). In the rest of the manuscript, we will describe how to express each hypothesis in a probabilistic form. Furthermore, we will describe a likelihood comparison approach for testing which hypothesis fits actual data, and whether different genomic regions conform to a single evolutionary model.

### Modeling dependencies of epigenomic changes on sequence mutations

#### Model N

Model N assumes that epigenomic changes are independent of both mutations and indels (Model N, Figure 2). We model evolutionary process of epigenomic changes as a Poisson process, in which the probability of change in time *t* is:

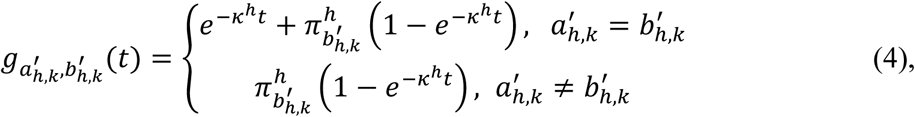

where 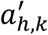 and 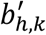 are binary states of the *h*^th^ histone modification on the *k*^th^ position of an alignment, and 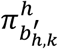 is the equilibrium probability of having the *h*^th^ histone modification on the *k*^th^ position, namely 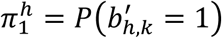 and 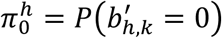. This probabilistic form is similar to the substitution model of DNA evolution [3]. On a position without indel 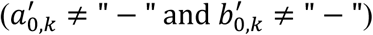, we have:

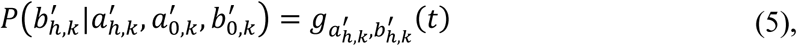

and

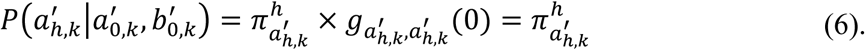

In order to model epigenomic changes on insertions and deletions, we introduce four parameters 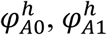 and 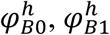 to represent local equilibrium probabilities of epigenomic states 0 and 1 in genomic regions *A* and *B*, respectively 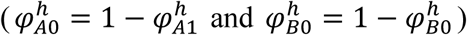. Unlike global equilibrium probabilities 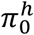 and 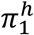, which are estimated from all homologous regions in the entire genomes, local equilibrium probabilities are estimated from each genomic region. On an insertion in the descendent sequence 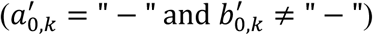, the transition probability is modeled as a mixture of the two transitions:

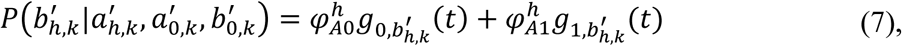

and because there is no place for histone mark on the *k*^th^ position in the ancestral sequence (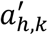 is not observed), we denote:

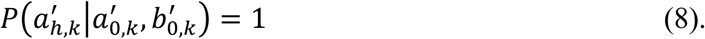

On a deletion in the descendent sequence 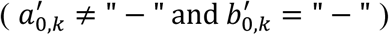, based on the reversibility of the evolutionary process we model the transition as a mixture of two transitions:

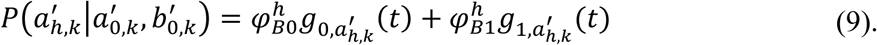

For the completeness of the model, we denote:

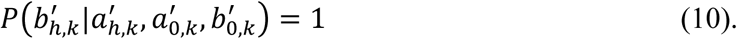

Taken together, 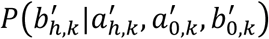 and 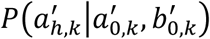 are given by:

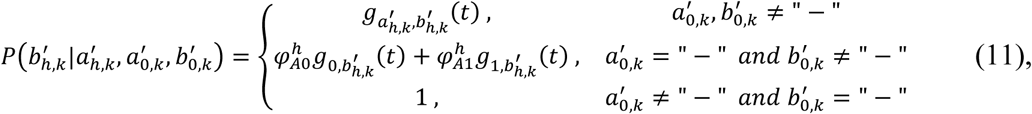

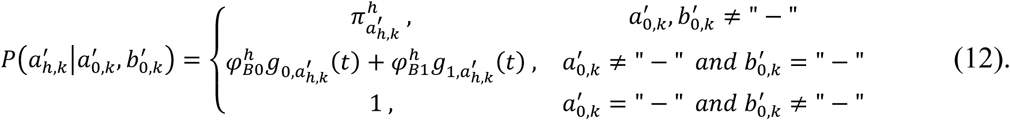

At this point, all the terms in the LCZ model have been specified. Equations (11) and (12) specify Model N, where epigenomic changes are independent of sequence changes.

#### Model M

In this model, epigenomic changes are dependent of sequence mutations but independent of indels (Model M, Figure 2). On a matched (no mutation) base at the *k*^th^ position in the alignment, namely 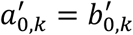, the epigenomic change is modeled with the Poisson process 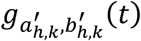 (Equation (4)), whereas on a position with mutation 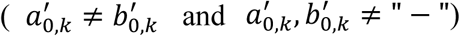, the descendent epigenomic state is modeled by global equilibrium probabilities,

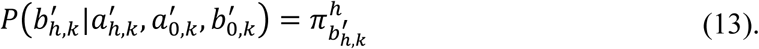

On indels, the epigenomic changes is modeled with the same approach as in Model N. Taken together, Model M is specified as:

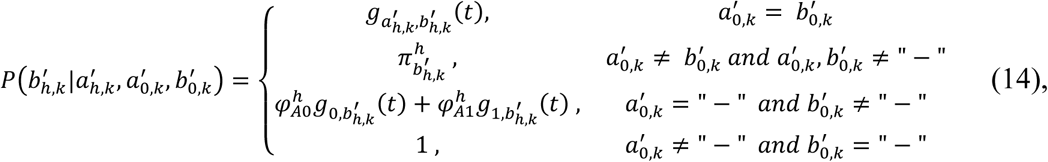

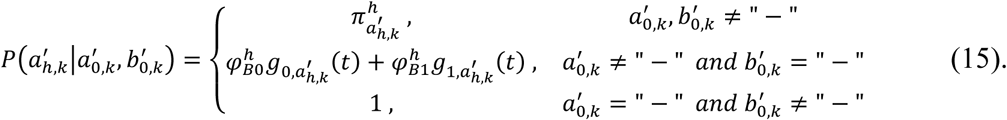

#### Model I

In this model, epigenomic changes depend only on sequence indels but not on mutations (Model I, Figure 2). On a position that is not indel 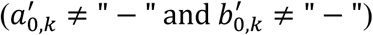, we use the same Poisson process (Equation (4)) as that in Model N to model the epigenomic changes.

On an insertion in the descendent sequence 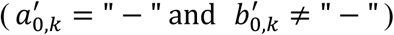, the transition probability becomes invariant of *t*:

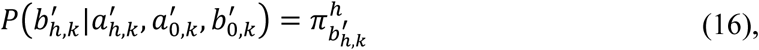

and since 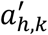 is not observed, we denote:

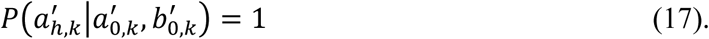

Similarly, on a deletion in the descendent sequence 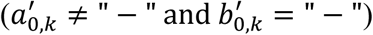, we have:

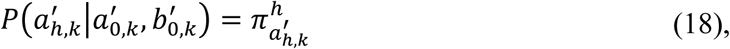

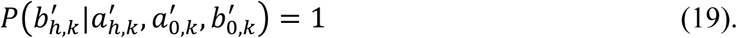

Altogether, 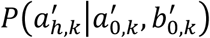 and 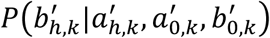 are given by:

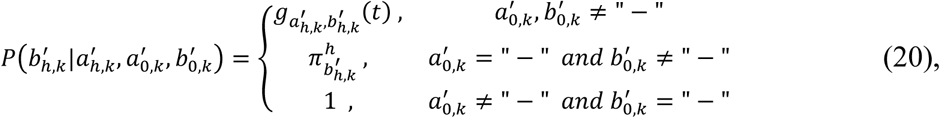

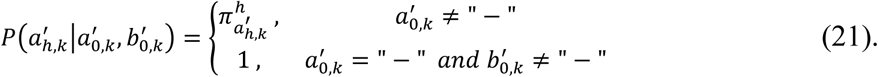

#### Model B

Model B assumes that epigenomic changes depend on both sequence mutations and indels (Model B, Figure 2). Similar to Model M, the epigenomic change is modeled with the Poisson process 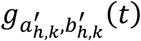 on a matched base 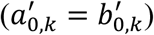, whereas on a position with mutation 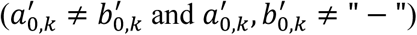, the descendent epigenomic state is modeled by the equilibrium probabilities. On indels 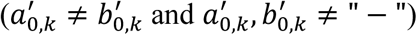, the epigenomic state is modeled using the equilibrium probability following Equations (16) – (19).

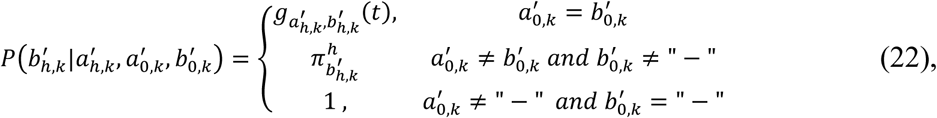

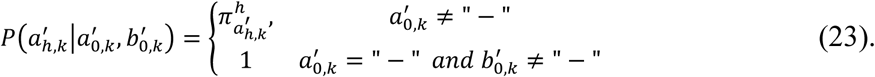

### A unified evolutionary model incorporating all four hypotheses

We express the probably of two homologous genomic regions as a mixture of the four models, thus obtained a general probabilistic model that do not depend on any of the specific hypothesis as follows:

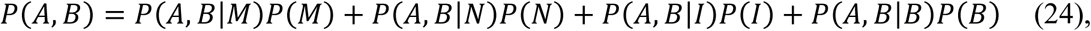

where *P*(*A*, *B*|*M*), *P*(*A*, *B*|*N*), *P*(*A*, *B*|*I*), and *P*(*A*, *B*|*B*) are the four probability density functions for Models M, N, I, and B, respectively.

### Development of a MLE algorithm for parameter estimation

We implemented a maximum likelihood estimation (MLE) algorithm for model fitting. The input data for the MLE algorithm are a list of pairs of homologous regions, hereafter termed *homologous pairs*, each of which contains two homologous sequences, and on each position of each sequence a binary indicator of state of each histone mark. The model parameters include equilibrium probabilities *π* and *φ*, birth and death rates *λ* and *μ*, substitution rate *s*, and the rate of change for each histone modification *κ*^*h*^. Our MLE calculation algorithm is a downhill simplex algorithm. The key for application of downhill simplex algorithm is being able to evaluate the likelihood function with given model parameters, which requires summing over all possible evolutionary paths between the two sequences, which was achieved by a dynamic program algorithm (Methods).

### Evaluation with simulation datasets

We tested performances of the models and the MLE algorithm with simulation data. First, we tested the convergence by comparing the estimated parameters at each iteration with the true parameters (Figure S1A). We simulated data with 8 sets of model parameters (Table S1, Methods) under each of the 4 models (Model M, N, B, I), resulting in a total of 32 datasets. Each dataset contained 100 pairs of 500bp-long homologous sequences and one histone modification on each sequence. We ran the MLE estimation algorithm twice with two initial values on each simulation dataset. Regardless of the initial values, the estimated parameters converged to true values in all simulated datasets (Figure S1A), and the negative log-likelihood function decreased monotonically (Figure S1B).

For a more comprehensive test, we simulated 10 datasets under each of the 4 models with each of the 8 sets of model parameters (Table S1), resulting in a total of 320 datasets. For each dataset we ran the MLE algorithm to convergence, and quantified the difference between the estimated parameters (*θ*) with true values (*θ*^∗^) with percent error (*e*), defined as *e* = (*θ* − *θ*^∗^)/*θ*^∗^. We summarized the percent errors from all the simulations for each true value (Figure S2). Regardless of the true values for *s*, *μ*, *κ*, the majority of the percent errors of all simulations were contained within 20% (|*e*| < 0.2). Greater variation of *e* were observed when the true values were very small (0.01). As the true values increased to 0.1 or 1, nearly all percent errors were contained within 10% (|*e*| < 0.1). We note that the estimated *κ* (rate of H3K4me3 switch) from real data was much larger than 0.1 (Table S2), and thus in the range where the estimated values nearly always converge to true values.

Next, we tested the capability of identifying the underlying model by comparison of likelihood functions. We generated 5 datasets (columns, Figure S3) under each hypothesis (Hypothesis M, N, I, or B, Figure S3), resulting in a total of 20 datasets. For each dataset, we computed the likelihood using every model (Model M, N, I, or B), resulting in four computed likelihoods (four dots in each column, Figure S3). In all simulation datasets, the model that resulted in the largest likelihood corresponded to the actual hypothesis from which the data were generated, suggesting that the true model corresponding the correct hypothesis could be identified by likelihood comparisons.

### Rates of sequence changes and H3K4me3 change between humans and rhesus monkeys

Our overriding question is whether interspecies changes of histone modifications depend on genomic sequence changes, and whether such dependence is invariant in the entire genome. Toward this goal, we used H3K4me3 changes in primate spermatids as a testbed system. We approached the above question with two major steps. First, we estimated sequence change rates and H3K4me3 change rate, and assessed the sensitivity of these estimates to model assumptions and to data processing procedure. We retrieved public epigenomic data from rhesus macaque and human in round spermatids (GSE68507) [28]. We estimated the sequence change rates (*s*, *μ*) and H3K4me3 change rate (*κ*) from each of the four models. We will not separately provide *λ* in results because *λ* is determined by homologous sequence lengths and *μ* [4]. Our estimation of *s*, *μ*, and *κ* were based on the union of H3K4me3 marked regions [28] and all DNase hypersensitive regions from 95 human cell lines [34], that was a total of 2,824,711 homologous genomic regions. The four models yielded nearly the same estimates for each parameter, where sequence substitution rate *s* was approximately 0.07, deletion rate *μ* was approximately 0.04, and H3K4me3 change rate *κ* was approximately 0.75 (Table S2). Executing the MLE algorithm 3 times with different initial values converged to nearly the same estimated values. These values are in line with the reports of large amounts of interspecies histone modification changes on homologous sequences, in the same cell type [35]. To assess the sensitivity of these estimates, we re-estimated the parameters with randomly sampled subsets of the homologous genomic regions (Table S2), and also with re-defined peak regions by applying different thresholds in ChIP-seq peak calling (Table S3). The estimated parameters by large were insensitive to these alternations, with an expected exception that *κ* exhibited a modest decrease when stringency for peaking calling drastically increased. This is because when few peaks were called from either species (q-value = 0.001, Table S3), the histone modification would not appear to change (no modification in either species).

### Epigenome-to-genome dependency in evolution is not uniform across the genome

Next, we compared the four evolutionary hypotheses on every homologous sequence pair and derived a genome-wide catalogue of the correspondence between genomic region and the best fit evolutionary model. Nearly the entire mappable portion of the human genome (effective genome) has homologous sequence in rhesus macaque genome. Approximately 5.5% of the homologous sequences were covered by H3K4me3 peaks in either species, accounting for 132,294 homologous pairs. For every pair, we computed the likelihood under each of the four models, and classified each homologous pair to one of the models according to the largest likelihood. A total of 73% of homologous pairs were classified to Model M, I, or B, where histone modification variation is dependent of DNA sequence changes (Figure 3A). The majority of these homologous pairs were classified to Model B, where histone modification variation is dependent of both sequence mutation and indel. On the other hand, a total of 27% of homologous pairs were classified with Model N, where histone modification variation does not depend on DNA sequence changes. These data are in line with an idea that the evolutionary changes of the genome may not completely determine all evolutionary changes of the epigenome.

**Figure 3.**
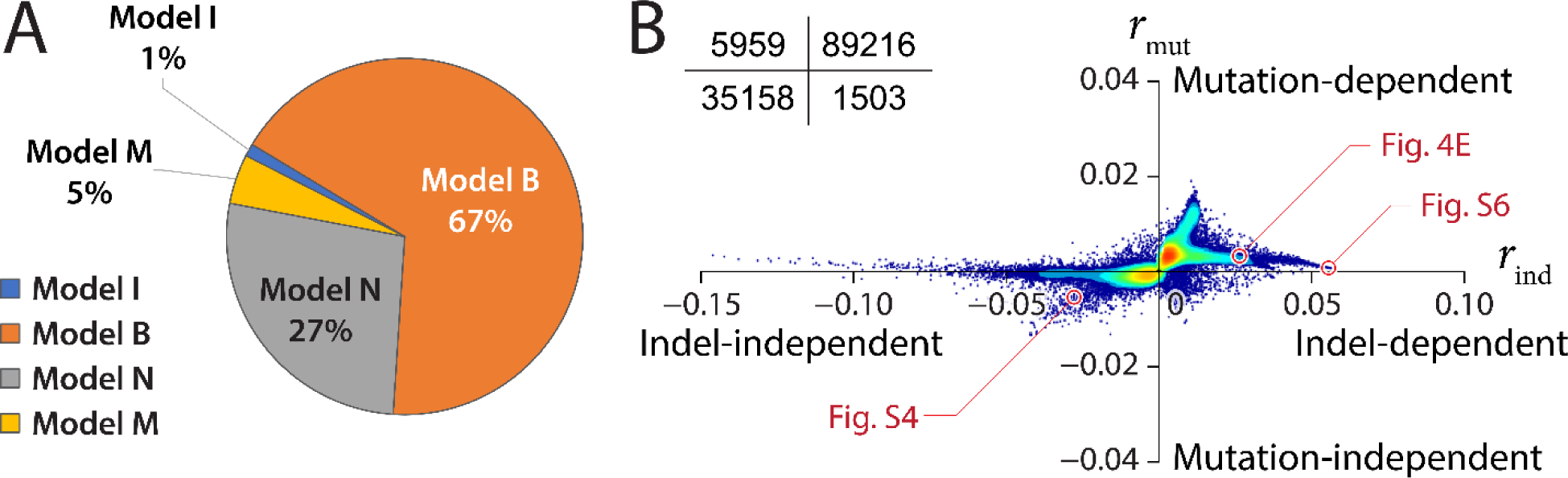
Classifications of homologous genomic regions into four models. (A) Proportions of human-macaque homologous regions classified into each model. (B) Scatterplot of all homologous regions showing the degree of dependence to indels (*r_ind_*, x axis) versus the degree of dependence to mutations (*r_mut_*, y axis). Actual data for selected homologous regions (red circles) are given in Figures 4E, S4, and S6. Insert: the numbers of homologous regions in each quadrant.

### Separating contributions of mutations and indels to epigenome-to-genome dependence

We asked whether sequence mutation or indel better accounts for epigenome-to-genome dependence in evolution. Toward this goal, we derived two metrics *r*_*mut*_ and *r*_*ind*_ to quantify the degrees of dependence of histone changes on mutations and on indels, respectively (Methods). These metrics were derived from a variation of likelihood-ratio test, where *r*_*mut*_ quantifies the overall fit of a homologous pair to Models N or I (independent of mutations) versus to Models M or B (mutation dependent), and *r*_*ind*_ quantifies the overall fit to Models N or M (independent of indels) versus to Models I or B (indel dependent). We quantified *r*_*mut*_ and *r*_*ind*_ for every homologous pair, and used a scatterplot to visualize the degrees of H3K4me3-to-mutation dependence (*r*_*mut*_, y axis) and H3K4me3-to-indel dependence (*r*_*ind*_, x axis, Figure 3B) of all the analyzed homologous pairs (132,294 in total). Overall, the homologous pairs exhibited greater variations of *r*_*ind*_ than *r*_*mut*_. The majority of homologous pairs exhibited *r*_*mut*_ close to 0, for example 110,400 (83%) homologous pairs exhibited |*r*_*mut*_| < 0.004. Data of these homologous pairs cannot clearly infer H3K4me3-to-mutation dependence. A greater number of homologous pairs exhibited non-zero *r*_*ind*_, including 8,965 homologous pairs with *r*_*ind*_ > 0.01, in which H3K4me3 changes are likely attributable to indels. Nearly no homologous pair exhibited H3K4me3 variation that is solely depend mutation (2^nd^ quadrant, Figure 3B), and in some homologous pairs neither mutation or indel appeared to relate to interspecies variation of H3K4me3 (3^rd^ quadrant in Figure 3B, Figure S4).

### Contribution of transposon induced indels to DNA-dependent H3K4me3 changes

We asked to whether indels induced by different transposon families exhibit similar impacts to interspecies variation of epigenome. To this end, we first classified species-specific transposon insertions into three groups, that are with no change to H3K4me3 (conserved peak), transposon insertion together with addition (transposon-induced peak) or removal (transposon-disrupted peak) of H3K4me3 (Figure 4A). Next, for each group we identified the number of contributing transposons from every transposon family.

**Figure 4.**
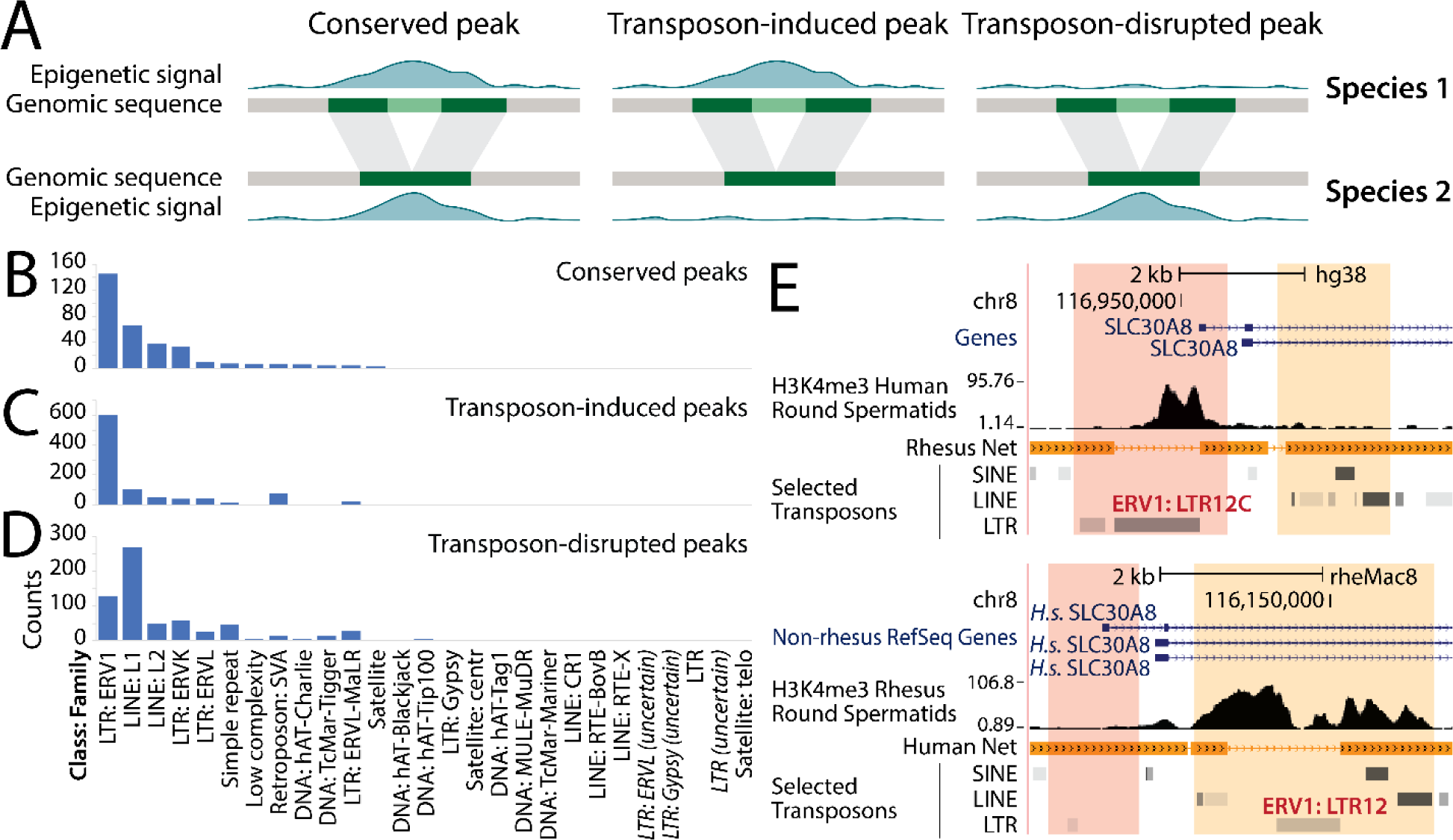
Classes of interspecies covariations of transposons and H3K4me3 peaks. (A) Three classes of covariations of transposon and H3K4me3 peaks. Shaded bands between two species indicate homologous sequences. Light green sequence: insertion of a transposon in Species 1. (B-D) Transposon copy number of each transposon family in conserved peaks (B), transposon-induced peaks (C) and transposon-disrupted peaks (D). (E) A homologous genomic region where interspecies variation of H3K4me3 peaks was associated with ERV1 insertions. Pink bands: a pair of homologous sequences in humans (upper panel) and macaque (lower panel), with a human-specific insertion (ERV1:LTR12C) as well as a human-specific H3K4me3 peak. Orange bands: another pair of homologous sequences, with a macaque-specific copy of ERV1 (ERV1:LTR12) and macaque-specific H3K4me3 peaks.

A total of 3,415 homologous pairs exhibited evidence (*r*_*ind*_ < −0.02) of H3K4me3 variation being independent of DNA changes, of which 330 (9.7%) contained species-specific transposons. Among these species-specific transposons that do not appear to interfere with H3K4me3, the endogenous retrovirus 1 (ERV1) family of long terminal repeats (LTR) was the most abundant transposon family, accounting for 143 (43%) of the conserved peaks (Figure 4B). This trend did not change when we altered the threshold into *r*_*ind*_ < −0.04 (Figure S5).

A total of 8,965 homologous pairs exhibited evidence (*r*_*ind*_ > 0.01) of DNA-dependent H3K4me3 changes, including 975 homologous pairs with transposon-induced peaks, and 631 homologous pairs with transposon-disrupted peaks. The ERV1 family was the most abundant transposon family with transposon-induced peaks, accounting for 598 (61%) of all transposon-induced peaks (Figure 4C). This trend did not change when we altered the threshold into *r*_*ind*_ > 0.025 (Figure S5). The promoter region of the *SLC30A8* gene is an example in case (Figure 4E). This promoter region harbors two homologous pairs, with one in the upstream regions of human and macaque transcription start sites (pink regions, Figure 4E) and the other in the downstream of TSSs in both species (orange regions, Figure 4E). An ERV1 transposon was inserted in the human upstream region, on which is a clear H3K4me4 peak, whereas the macaque upstream region did not contain the ERV1 sequence and did not exhibit H3K4me3 (pink regions, Figure 4E). Furthermore, another ERV1 sequence was inserted in the downstream region in macaque, where H3K4me3 was installed, whereas the human homologous sequence did not have the ERV1 sequence and did not harbor any H3K4me3 peak (orange regions, Figure 4E).

Unlike transposon-induced peaks that were primarily concentrated to ERV1, transposon-disrupted peaks were contributed from a larger variety of transposons, including L1, ERV1, ERVK, simple repeat, L2, and ERVL-MaLR (Figure 4D). The L1 family was most abundant in this group, accounting for 265 (42%) of all transposon-disrupted peaks (Figure 4D). This trend did not change when we altered the threshold to *r*_*ind*_ > 0.025 (Figure S5). A case in point is at the *ZNF630-AS1* promoter, where a L1 transposon (a, Figure S6) was inserted specifically in the macaque promoter between two ERVL-MaLR family repeats (b, c, Figure S6), which was coupled with disappearance of H3K4me3. In summary, ERV1, L1, L2, ERVK, ERVL, simple repeat, SVA, and ERVL-MaLR are the most abundant transposons near H3K4me3 peaks. Transposon-induced peaks were most strongly associated with, and transposon-disrupted peaks were most strongly associated with L1 transposons.

## Discussion

In parallel to the plethora of evidence on DNA-dependent installation and removal of histone modifications, a smaller but increasing amount of data suggest trans-generation DNA-independent inheritance of histone modifications [32, 33]. It remains unclear how many generations could DNA-independent epigenetic inheritance endure, or more importantly whether it is preserved in evolutionary timescale. By initiating probabilistic models of epigenome-genome evolution, this work begins to offer a quantitative framework to address the above question. Future developments of epigenome-genome evolution models may begin to address questions including whether any evolutionary selection acts on the epigenome independently of the genome, and whether any selection forces were received jointly by genome and epigenome. Therefore, we anticipate integrated analyses of genome-epigenome data to expand the domain of evolutionary biology, and the development and deployment of epigenome-genome evolution models to be essential for this expansion.

We foresee a number of improvements to this initial attempt of modeling. First, in this work we have only considered binary states of histone modifications. To remove this assumption, 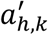 and 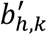 can be allowed to take any finite discrete numbers, in which case the form of Equation (4) does not change and hence the forms of the rest of the models do not change. Second, the conditional independence assumption can be removed. To model the dependent changes of two histone modifications, for example H3K4me2 and H3K4me3, the two modifications can be coded with the same index (*h*) and let 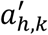 and 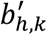 to take the following form:

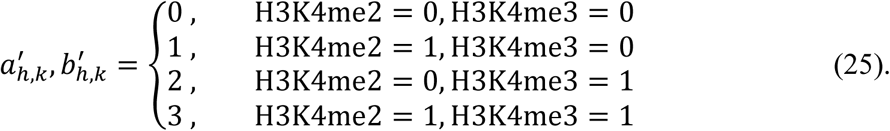

Transposon insertions were associated with gains and losses of H3K4me3 peaks. While H3K4me3 losses were associated with a variety of transposon families, H3K4me3 gains were enriched with the ERV1 family of transposons. The latter may be a result of species-specific recruitment of transcription factors. In line with this idea, ERV1 was the most notable transposon family involved in species-specific binding of pluripotency regulators OCT4 and NANOG in embryonic stem cells [36]. Our *de novo* motif search revealed a total of 31 DNA motifs that were enriched in ERV1 transposons as compared to other LTRs (Homer p-value < 10^−40^), where the most significant motifs resembled the binding motifs of NFYB (a.k.a. CCAAT box, Homer p-value < 10^−94^), HOXC13 (p-value < 10^−90^), BARX1 (p-value < 10^−89^), and LIN28A (p-value < 10^−84^). According to gene expression data of 37 human tissues from Genotype-Tissue Expression (GTEx) [37] and Human BodyMap 2.0 that were normalized and visualized by Genecards (www.genecards.org), Nfyb was expressed in nearly all human tissues, whereas Hoxc13, Barx1, and Lin28a were all most strongly expressed in testis. Lin28a exhibited 10 times greater expression in testis than in any other analyzed human tissues. The CCAAT box is capable of recruiting ASH2L, a component of the MLL histone methyltransferase complex responsible for H3K4 methylation [38]. These data suggest a model for ERV1 mediated induction of species-specific H3K4me3 in spermatids. ERV1 harbors binding motifs of testis-expressed transcription factors as well as the CCAAT box. Species-specific ERV1 sequences recruit testis-induced HOXC13, BARX1, LIN28A that help to recruit NFYB and the MLL complex, which in turn establish species-specific H3K4me3 peaks (Figure S7). Finally, the human-specific and macaque-specific insertions of two copies of ERV1 appeared to have induced H3K4me3 in respective insertion regions, near the Slc30a8 promoter in both species (Figure 4E), providing a potential example of convergent evolution mediated by species-specific transposon insertions.

## Methods

### Maximum likelihood estimates

The MLE of *π* and *φ* are calculated by frequency estimates. Due to the relationship 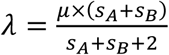 [4], the MLE of *λ* is determined as long as the MLE of *μ* is determined. Evolutionary time *t* is set to 1. The remaining parameters *μ*, *s*, and *k*, collectively denoted as *θ*, are obtained by minimizing the negative log-likelihood function,

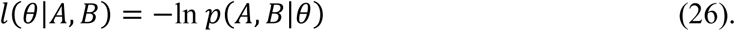

We used simplex downhill algorithm for this optimization. In each iteration, a number of new parameter sets *θ*^new^ were generated. The algorithm iteratively evaluated each *l*(*θ*^new^|*A*, *B*) and compared it with *l*(*θ*^old^|*A*, *B*) to select the best one, until the minimum was achieved. We developed a dynamic program algorithm to evaluate *l*(*θ*|*A*, *B*) for each model.

### Dynamic programming algorithms

#### Model I and Model B

The dynamic programming algorithm was implemented following the idea described in [39], which was a simplification of the procedure described in [4]. This procedure has been shown to vastly reduce the runtime of the algorithm. Since *P*(*A*, *B*) = *P*(*A*)*P*(*B*|*A*), and *P*(*A*) can be directly computed, only the computation of *P* (*B*|*A*) requires dynamic programming. Denote the (*s*_*A*_ + 1) × (*s*_*B*_ + 1) matrix in the dynamic programming algorithm by *L*, *L* was computed following rules listed below.

##### Boundary conditions

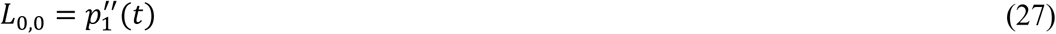

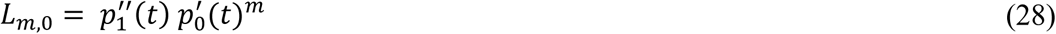

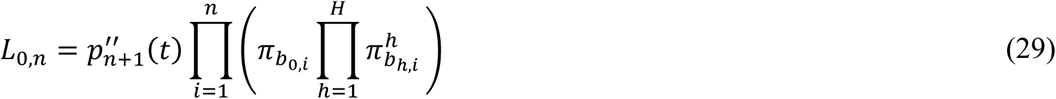

##### Recursive procedure

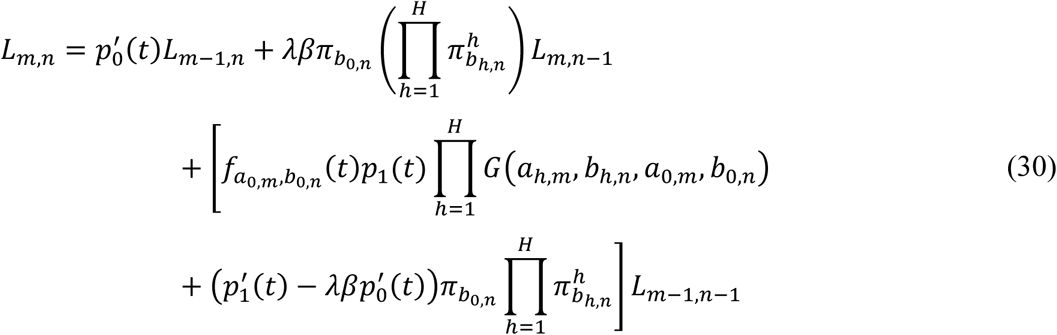

All of the functions *p*, *p*′, *p*″ are given by [4]. For Model I, the term *G*(*a*_*h*,*m*_, *b*_*h*,*n*_, *a*_0,*m*_, *b*_0,*n*_) is *g_a_h,m__,_b_h,n__* (*t*). For Model B, the term is the same as for Model I on matched bases, while it becomes 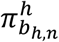 on mismatches.

#### Model N and Model M

The dynamic programming algorithm was implemented using a similar idea as for Model I and B, except that the computation of *P*(*A*^*h*^) became part of the recursive procedure, and the function *G* changed. Given equations (13) and (14) in the main text:

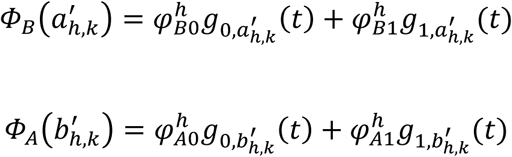

The matrix *L* was computed following rules listed below.

##### Boundary conditions

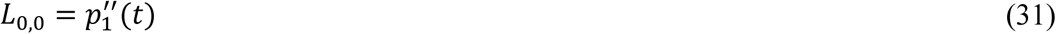

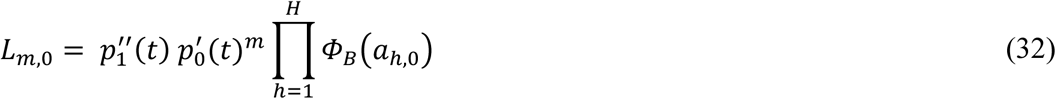

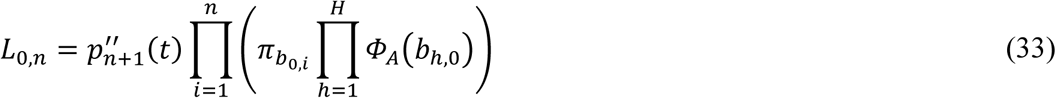

##### Recursive procedure

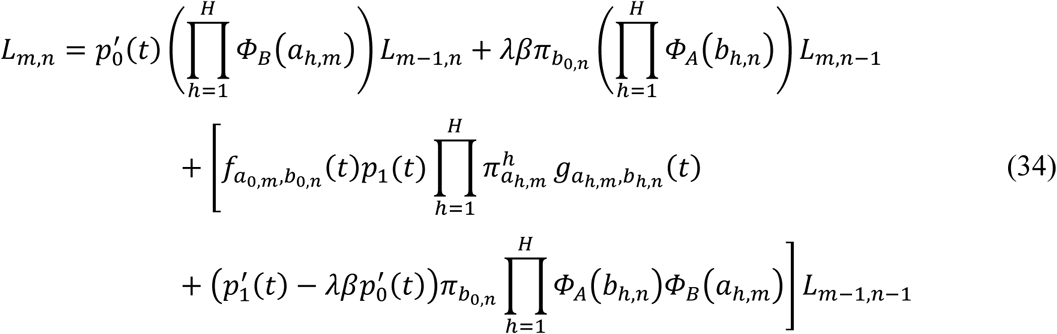

The term *G*(*a*_*h*,*m*_, *b*_*h*,*n*_, *a*_0,*m*_, *b*_0,*n*_) is *g_a_h,m_,b_h,n__* (*t*) for Model N (same as Model I). The term for Model M is the same as for Model N on matched bases, while it becomes 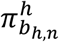 on mismatches (same as Model B).

### Generating simulation datasets

In the simulation test, we used only one histone modification. The equilibrium probabilities were set to *π*_*A*_ = *π*_*C*_ = *π*_*G*_ = *π*_T_ = 0.25, *π*_0_ = 0.9, *π*_1_ = 0.1. The simulation data contains 100 500-base-long sequence pairs unless otherwise specified.

#### Model I and Model B

The simulation data was generated based on the corresponding hypotheses of models. The ancestor was first generated by drawing bases and histone modification states randomly based on the equilibrium probabilities. For each link in the ancestral sequence, the number of its descendent link was drawn from the following distribution:

##### Immortal link

Assume that random variable *n* represents the number of descendent links of an immortal link. *n* follows the distribution:

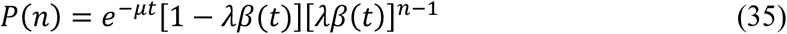

##### Normal link

Assume that the random variable *n* represents the number of descendent links of a normal link. *n* follows the distribution:

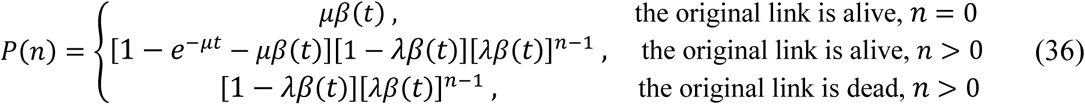

where 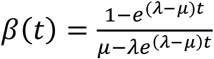.

On insertions, the bases and histone modification states were drawn based on the equilibrium probabilities. On matches, the bases and histone modification states were determined based on the substitution probabilities. On mismatches, the bases and histone modification states were also determined based on the substitution probabilities for Model I, while they were drawn based on the equilibrium probabilities for Model B.

#### Model N and Model M

For each region pair, the ancestral sequence was first generated by drawing bases randomly based on the equilibrium probabilities. Local equilibrium probabilities 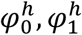 were drawn from a beta distribution 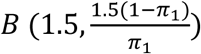 and assigned to the ancestral region. After that, the path was determined using equation (35) and (36). On insertions, the descendent nucleotides were drawn based on the equilibrium probabilities, and the histone modification states were drawn based on the local equilibrium probabilities. On matches, the bases and histone modification states were determined based on the substitution probabilities. On mismatches, the bases and histone modification states were also determined based on the substitution probabilities for Model N, while they were drawn based on the equilibrium probabilities for Model M.

### Analysis of human and rhesus macaque H3K4me3 ChIP-Seq data

We first fitted the four models and obtained MLE from each model. Our training set consisted of a set of homologous region pairs that were potentially marked by H3K4me3. These regions were identified by merging three types of regions: (1) The DNase I hypersensitivity peak clusters derived from 95 human cell lines[34], as cis-regulatory elements marked by histone modifications are usually hypersensitive to DNase I; (2) H3K4me3 peak regions identified from the human RS H3K4me3 ChIP-Seq data using MACS2 [40]; (3) H3K4me3 peak regions identified from the macaque RS H3K4me3 ChIP-Seq data. The merged regions were remapped to rhesus macaque’s genome using liftOver [41] to find their homologous regions, which yielded 2,824,711 region pairs. After that, 8,000 region pairs were randomly sampled for parameter estimation to reduce the runtime of our MLE program. We always started from two initial guesses to avoid local minima. All estimated parameters converged to the same values regardless of initial guesses, showing that the global minima of the negative log-likelihood functions were achieved. We randomly sampled the data for three times, and the estimated parameters remained nearly the same with coefficient of variation smaller than 0.05, showing that the data sampling didn’t lead to biases to estimated parameters (Table S2).

To assess to what extent the MLE was sensitive to peak calling results, we also conducted parameter estimation with Model I and B using two other q-value cutoffs for MACS2 peak calling. None of the parameters changed vastly as the q-value changed, showing that the parameter estimation process was not very sensitive to peak calling results (Table S3).

### ChIP-Seq data pre-processing

ChIP-Seq datasets were mapped to human genome assembly hg38 and rhesus macaque genome assembly rheMac8 using bowtie2 with default settings. Duplicated reads and reads with MAPQ<6 were then removed from the data. Peaks were identified using MACS2 with the “broadpeak” option.

### Sensitivity test of MACS2 q-value threshold

We used three different q-value thresholds when using MACS2 to identify peaks to test the sensitivity of our models to MACS2 q-values. As the q-value cutoff decreased, fewer peaks were detected from the data, leading to decreased equilibrium probability *π*_1_. Among the three parameters, *s* and *μ* remained identical regardless of the cutoff used, as the two parameters described changes of DNA sequences and wouldn’t change as long as the sequences were unaltered. On the other hand, *κ* decreased slightly as the cutoff decreased, which was because the data became more conserved when fewer peaks were identified. The results are shown in Table S3.

### Generating candidate regions for parameter estimation and model competition

Peak regions identified in rhesus macaque RS H3K4me3 ChIP-Seq data were first remapped to human genome using liftOver with minMatch=0.5. For parameter estimation, three files: (1) The DNase I hypersensitivity peak clusters derived from 95 human cell lines; (2) H4K3me3 peaks identified in the human RS H3K4me3 ChIP-Seq data; (3) remapped H3K4me3 peaks identified in the macaque RS H3K4me3 ChIP-Seq data were merged. The merge regions were trimmed to no longer than 500bp, and remapped to rhesus macaque genome using liftOver with minMatch=0.5 to find their homologous regions. After that, the H3K4me3 ChIP-Seq data was distributed to these region pairs based on the identified peak regions. Finally, 8,000 regions were randomly sampled for parameter estimation.

For region classification, only human and rhesus H3K4me3 peaks were merged. The merged regions were trimmed to no longer than 2,000bp, and remapped to rhesus genome using liftOver with minMatch=0.1. Remapped regions with less than 90% realigned successfully were extended to the length of the original ones. The H3K4me3 ChIP-Seq data was then distributed to these region pairs based on the identified peak regions.

### Classification of H3K4me3 peak regions in human and rhesus macaque

With the estimated parameters, we focused on H3K4me3 peak regions identified from human and rhesus macaque RS ChIP-Seq data to test the effects of sequence changes on epigenomic changes. Among all sequences in the human genome with homologous sequences in rhesus macaque, around 5.5% showed H3K4me3 in either species. These peak regions were merged together, trimmed to no longer than 2,000bp, and remapped to find their homologous regions, which yielded 132,294 region pairs. For each homologous pair, four likelihoods were obtained using the four models, leading to a 132,294 by 4 likelihood matrix. Each homologous pair was categorized to one model by the largest likelihood.

### Separating evolutionary impacts of sequence mutations and indels

We leveraged the four models to evaluate the relative impacts on interspecies epigenomic variations from mutations and that from indels. By integrating out mutation’s impacts from the probabilities models, we obtained the overall impacts of indels, and vice versa. We introduce the binary variable *mut* to indicate independence *mut* = 0 and dependence *mut* = 1 to mutations, and binary variable *ind* to indicate independence of dependence to indels. The probability of observing a region pair given *mut* or *ind* can be expressed as a combination of likelihoods yielded by different models:

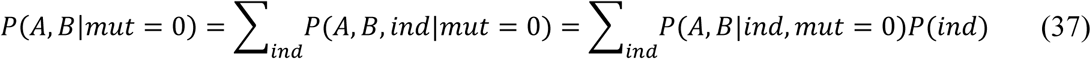

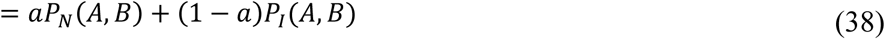

The coefficient *a* was estimated using the frequency of regions within which epigenomic changes are independent to indels. Similarly, we also have

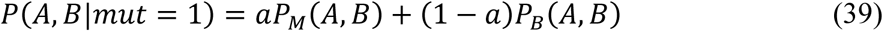

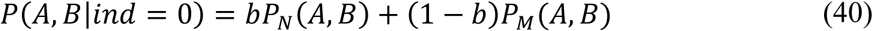

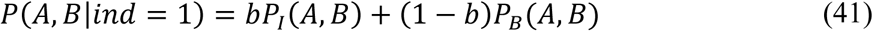

The coefficient *b* was estimated using the frequency of regions within which epigenomic changes are independent to mutations.

We proposed two normalized likelihood ratios, *r_ind_* and *r_mut_* to assess the effect of indels and mutations:

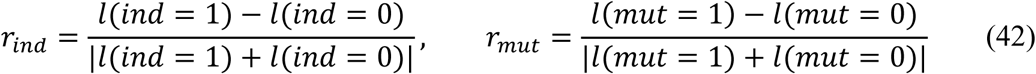

where *l*(*ind*) = log *P*(*A*, *B*|*ind*), *l*(*mut*) = log *P* (*A*, *B*|*mut*). *r*_*ind*_ and *r*_*mut*_ represented the dependency of epigenomic changes on indels and mutations. The extent of independence increased as the ratios increased.

### Searching for species-specific transposon insertions

We first selected 789 indel-independent regions with *r*_*ind*_ > 0.04 and 1,441 indel-dependent regions with *r*_*ind*_ < −0.025 for transposon analysis. We downloaded RepeatMasker files for hg38 and rheMac8 as transposon annotations. For each region pair, transposons longer than 500bp within the two regions were compared. Transposons with the same family, class and name were removed to keep species-specific transposon insertions. Regions with either human-specific or macaque-specific transposon insertions were kept and classified into three categories: conserved peak regions, transposon-induced peak regions and transposon-disrupted peak regions. Families and classes of species-specific transposon insertions were then summarized in each category. This analysis was repeated with 3,416 indel-independent regions with *r*_*ind*_ > 0.02 and 8,966 indel-dependent regions with *r*_*ind*_ < −0.01.

### Motif analysis

Sequence motifs were identified within 589 LTR-ERV1 transposons found in the 598 transposon-induced peaks with *r*_*ind*_ > 0.01 (target transposons). All LTR transposons in the ERV1 family longer than 500bp were used as background (background transposons). The *De novo* motif discovery was performed using Homer [39] with default parameters.

## Declarations

### Ethics approval and consent to participate

Not applicable.

### Consent for publication

Not applicable.

### Availability of data and materials

The datasets analyzed during the current study are available in the Gene Expression Omnibus (GEO accession number GSM1673960, GSM1673962, GSM1673983, GSM1673985) [28]. https://www.ncbi.nlm.nih.gov/geo/query/acc.cgi?acc=GSE68507 Codes for this work can be found at: https://github.com/Zhong-Lab-UCSD/EpiEvolutionaryModel

### Authors’ Contributions

All authors developed the models. SZ conceived the project. JL developed algorithms, conducted simulation tests and data analysis. All authors wrote the manuscript.

## Acknowledgement

We thank Dr. Dan Xie and Xin He for valuable discussions. This project is funded by NIH R01HG008135, DP1HD087990.

